# Serological evidence of SARS-CoV-2 infection in red and fallow deer in Great Britain

**DOI:** 10.1101/2025.10.17.682775

**Authors:** Joanna Vieira O’Neill, Joash Tan, Ed Long, Charlotte L. George, Joey Olivier, Chloe Qingzhou Huang, Hazel Stewart, Diego Cantoni, Chris Davis, Kristin Waeber, Connor G. G. Bamford, Craig P. Thompson, Nigel J. Temperton, Jonathan L. Heeney, George W. Carnell

## Abstract

SARS-CoV-2 is the viral agent of COVID-19 disease in humans and has been shown to infect a wide range of mammals, including white-tailed deer in North America and fallow deer in the Republic of Ireland. There are six species of deer in the UK which inhabit both urban and rural areas, providing a broad interface for human-deer interaction. Little is known if British deer species act as a reservoir for zoonotic spread of consequence to human health. Here, we report a high seroprevalence of SARS-CoV-2 neutralising antibodies in British deer, adding native red deer to the list of susceptible species. 62% of fallow and 25% of red deer sampled between 2024 and 2025 from one British deer park were positive for SARS-CoV-2 infection by serology. Data suggest that deer may function as permissive, reservoir-capable hosts and highlight the risks of anthroponosis and zoonosis from unregulated human-deer contact. While there is currently no evidence to suggest that these viruses are persisting within deer populations in Great Britain, the animal origins of SARS-CoV-2 highlight the importance of appropriate and limited interactions with wildlife, to minimise transmission risks and safeguard both animal and human health.

## INTRODUCTION

Severe acute respiratory disease syndrome coronavirus-2 (SARS-CoV-2) spilled over from wildlife to humans in late 2019, resulting in a highly transmissible acute respiratory disease in humans and leading to the global “COVID-19” pandemic^1,2^. The virus continues to circulate in humans to this day^3^. Much work has been carried out on the entry dynamics of this virus, identifying the factors that facilitated its spillover into humans, rapid establishment, circulation, and the multiple evolutionary steps leading to SARS-CoV-2 becoming endemic.

The prototype “Wuhan” strain of SARS-CoV-2 rapidly underwent evolution in humans, giving rise to new variants of concern (VOCs) starting with Alpha (B.1.1.7). Each VOC was sequentially replaced by new VOCs with evolutionary advantages. Alpha was replaced with Beta (B.1.351), Gamma (P.1), Delta (B.1.617.2), and then Omicron (B.1.1.529). First detected in South Africa and Botswana in November 2021, Omicron was characterised by increased transmissibility and immune evasion^4–7^. Further Omicron sub-variants emerged and some underwent subsequent recombination, producing a multitude of Omicron sub-lineages, which have continued to dominate until the present day^8,9^.

Angiotensin-converting enzyme 2 (ACE2) is a host receptor and key determinant of species tropism for SARS-CoV-2^10^. The high homology of ACE2 and the subsequent broad host tropism which SARS-CoV-2 displays, means that the virus has been detected in mammals worldwide in parallel with the human pandemic^11–17^. Among the multitude of reported susceptible mammalian hosts, SARS-CoV-2 was documented to have infected white-tailed deer populations in the USA and Canada, including genomic evidence for within-herd circulation and transmission to neighbouring human populations^18–24^.

Wild deer are present throughout the United Kingdom, with an estimated population of up to two million, including captive and enclosed deer found in deer parks, farms, and other zoological collections^25^. There are six species of wild deer in Great Britain, of which two are native (red and roe), whilst fallow, sika, muntjac and Chinese water deer are introduced species. High-risk factors for disease transmission in British deer ecology include: sympatric multi-species deer populations, free movement of wild deer, interactions with domestic livestock, and rapid growth of deer populations^26^.

ACE2 is expressed in cells of the lower respiratory tract of deer species found in Great Britain including red deer, roe deer and fallow deer, identifying these deer species as potential susceptible hosts^27^. SARS-CoV-2 has been reported in fallow deer in a deer park in Dublin, representing the only positive study resulting from samples taken from a managed deer park of high human use, and indicating a broad human/deer interface exists and there is significant potential for interaction and viral spillover^28^.

Several coronaviruses are endemic in both human and animal populations. Human coronaviruses (HCoVs) OC43, HKU1, 229E, and NL63 are endemic in humans and are believed to have originated from animal reservoirs, typically causing self-limiting, mild respiratory tract infections in people^29^. HCoV-OC43, in particular, is believed to have arisen through zoonotic transmission of a bovine coronavirus (BCoV), reinforcing the interconnectedness of human and animal viral populations^30^.

BCoV is a *Betacoronavirus*, belonging to the same genus as SARS-CoV and SARS-CoV-2, and causes both enteric and respiratory disease in cattle. Furthermore, BCoV and bovine-like coronaviruses are able to infect white-tailed deer and other wild ruminants^31,32^. These findings support the targeted surveillance of wild and domestic animal populations known to harbour coronaviruses of potential significance to human health.

Here, we report SARS-CoV-2 seropositivity in a single British deer park with red deer (*Cervus elaphus*) and fallow deer (*Dama dama*), across three cohorts culled between February/March 2024 and February/March 2025. These deer were culled during routine population management, enabling sample collection for this study without affecting which animals were selected for culling. From a total of 332 animals, we report a 43% positive rate (62% in fallow deer, 25% in red deer) to a panel of diverse SARS-CoV-2 variants, including contemporary Omicron-lineage strains such as BA.2.86 and JN.1^33,34^. These data are a strong indicator for significant spill over from humans to deer, adding complexity to SARS-CoV-2 disease ecology in Great Britain.

Ethical approval was granted by the Animal Welfare and Ethical Review Board, University of Cambridge. No identifying information has been included for the deer or regions where samples were taken. These will be published after peer-review.

## RESULTS

Sera were collected at three different time points from a total of 332 deer culled in February/March (spring) 2024, November/December (autumn) 2024, and February/March (spring) 2025. The spike-bearing lentiviral pseudotype virus (PV) system was used to assay neutralising antibodies in the deer sera^35^. A broad panel of spike proteins were used, which arose in humans at various times: 2019 (Wuhan), 2021 (Omicron sub-variant BA.1), 2022 (Omicron sub-variant BA.2.75), 2023 (Omicron sub-variant XBB.1.5) and 2024/2025 (Omicron sub-variants BA.2.86, JN.1). Seropositivity was defined as a half maximal inhibitory concentration (IC_50_) of neutralising antibody above the sensitivity cutoff of the assay (Log_10_40), with PV-only and cell-only controls used to normalise the assay neutralisation.

### Demographics

Of 332 deer serum samples, 174 (52%) were from red deer and 158 (48%) were from fallow deer (Table 1).

**Table 1.**
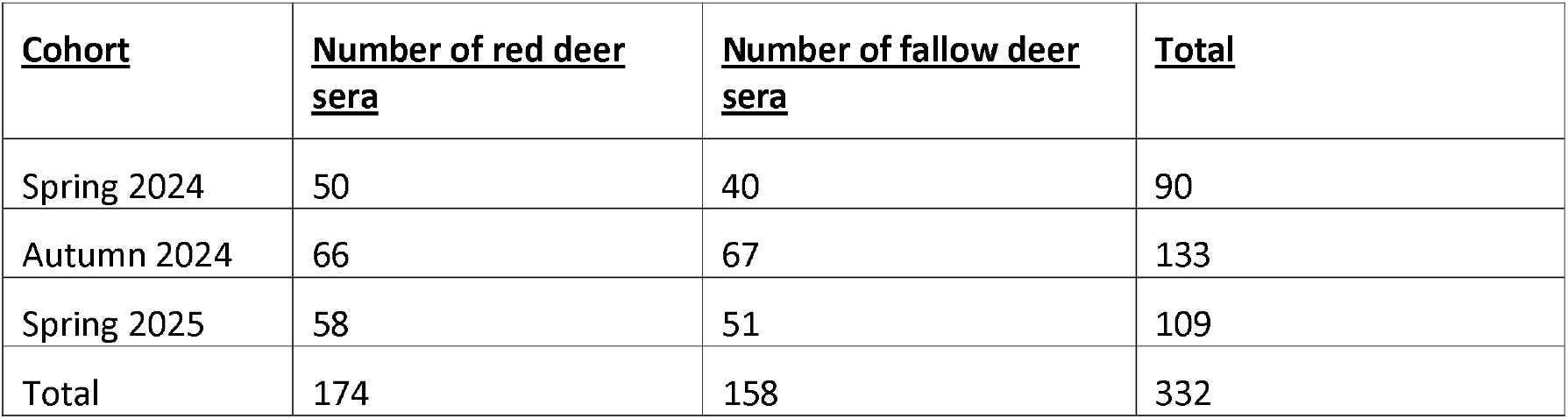
Species demographics of the 332 deer samples.

Across the three cohorts, 197 (59%) deer were male and 135 (41%) were female (Table 2). Male deer were culled in the spring cohorts and female deer were culled in the autumn cohort, in order to maintain healthy breeding populations. Our samples were primarily from the 2Y age group, which includes yearling deer up to and including 2-year olds. The fewest samples were obtained from calves under 1 year of age (Table 2). The median age of each cohort was 2 years old in spring 2024, and 3 years old in autumn 2024 and spring 2025. Across the three cohorts, the youngest deer culled were 1 month old and the oldest deer was 11 years old.

**Table 2.** Age and sex demographics of each cohort. Nb. Age data was unavailable for one deer sample.

| <u>Cohort</u> | <u>Age group</u> |  |  |  |  |  | <u>Sex</u> |  |
| --- | --- | --- | --- | --- | --- | --- | --- | --- |
|  | ≤ 1 year old (≤1Y) | 2 years old (2Y) | 3 years old (3Y) | 4 years old (4Y) | 5 years old (5Y) | > 5 years old (>5Y) | Male | Female |
| Spring 2024 | 1 | 48 | 15 | 8 | 8 | 10 | 90 | 0 |
| Autumn 2024 | 5 | 50 | 36 | 16 | 12 | 13 | 4 | 129 |
| Spring 2025 | 5 | 41 | 30 | 11 | 11 | 11 | 103 | 6 |
| <u>Total</u> | 11 | 139 | 81 | 35 | 31 | 34 | 197 | 135 |

### Neutralising antibody seroprevalence

Neutralising antibody responses against at least one SARS-CoV-2 spike variant PV were detected in 43% (142/332) of all deer samples across the three cohorts, as defined by a neutralising antibody IC_50_ value above our lower assay cutoff. Seroprevalence was lowest against the ancestral Wuhan spike which circulated in 2019/2020, whereas substantially higher responses were observed to more recent Omicron-lineage variants (Figure 1). Seropositivity rates were highest against the XBB.1.5 spike which predominated in 2023, at 33% (108/332) of all samples (Figure 1). By comparison, seroprevalence to more recently circulating lineages BA.2.86 and JN.1 PVs were lower, at 18% (61/332) and 20% (66/332) of all samples, respectively (Figure 1). These findings indicate that recent exposures within the deer population were predominantly associated with Omicron-lineage variants, such as XBB.1.5, rather than with the ancestral Wuhan strain.

**Figure 1.**
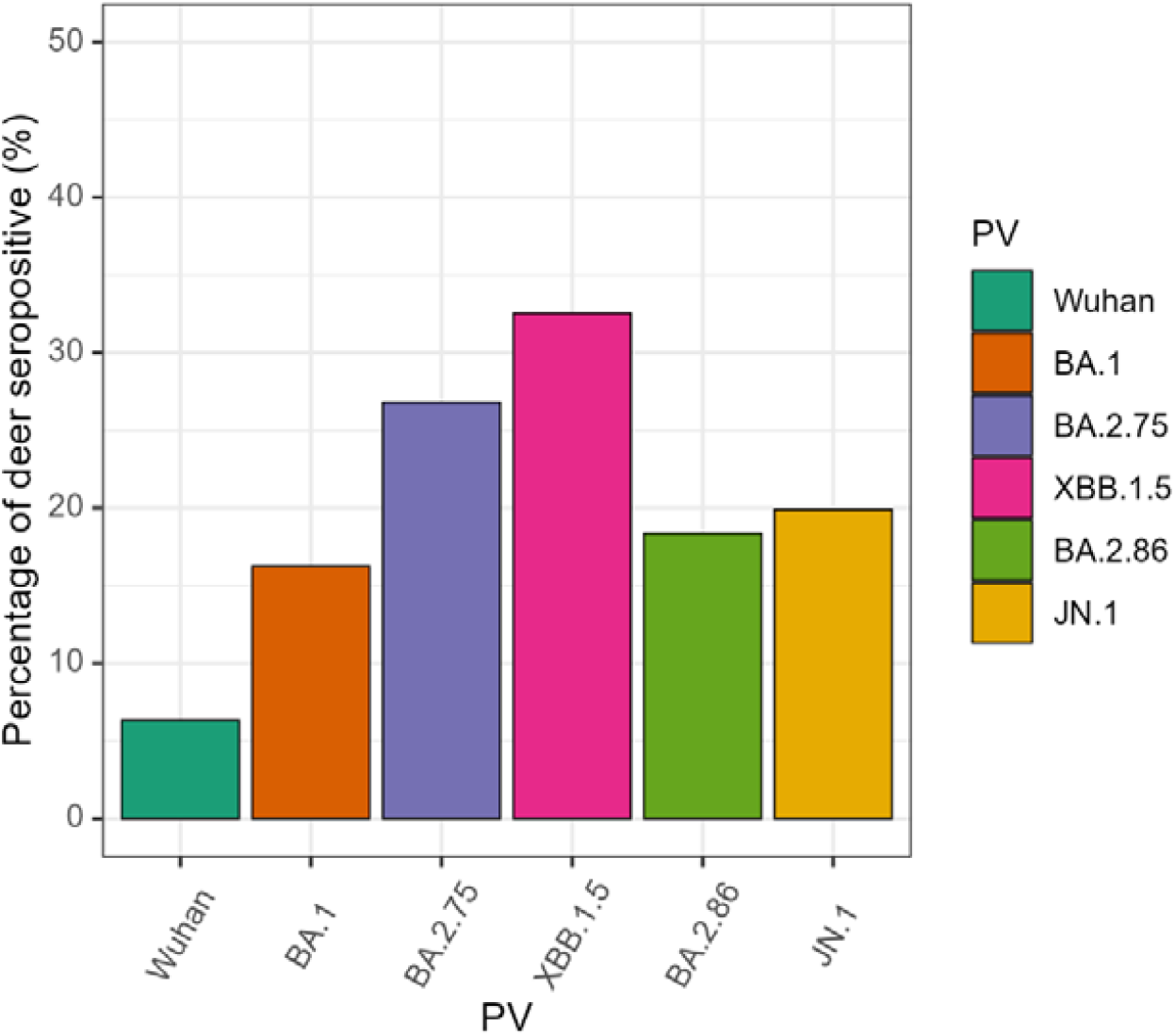
Seroprevalence of positive neutralising antibody responses to Omicron spike-bearing lentiviral pseudotypes (PVs): Wuhan (2019), Omicron sub-variants BA.1 (2021), BA.2.75 (2022), XBB.1.5 (2023) and BA.2.86 and JN.1 (2024/5). Seropositivity defined by IC_50_ values above the assay lower cutoff.

### Temporal trends

The highest seropositivity rates were observed in early spring 2024, and there was a decrease in seropositivity with subsequent cohorts. 47% (42/90) of the spring 2024 cohort were seropositive for neutralising antibodies to at least one SARS-CoV-2 spike PV, compared to 41% (55/133) seroprevalence in autumn 2024 and 41% (45/109) in spring 2025 (Figure 2).

**Figure 2.**
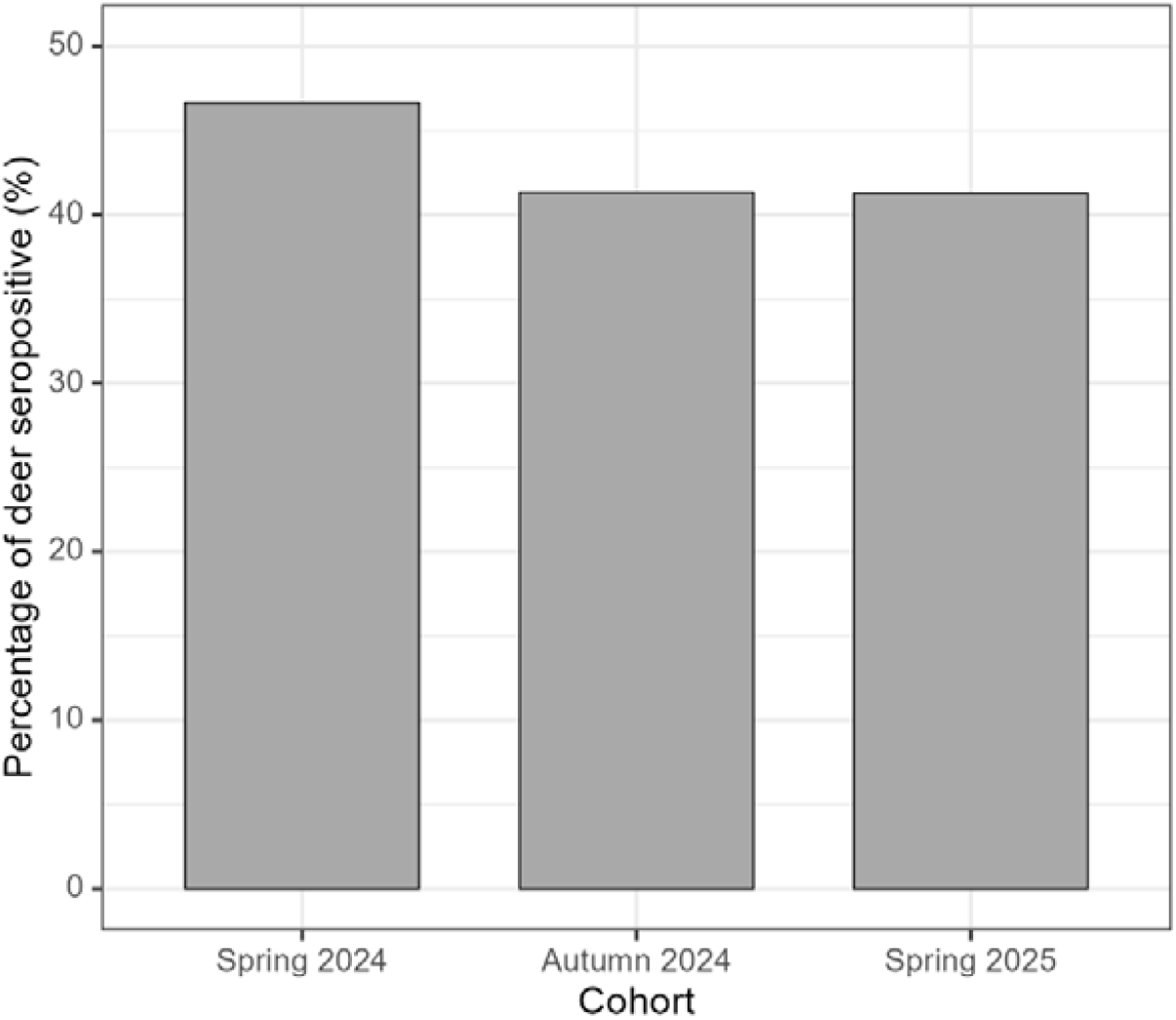
Seroprevalence of positive neutralising antibody responses to at least one SARS-CoV-2 spike variant PVs. Responses stratified by each cull cohort: spring 2024, autumn 2024 and spring 2025. Seropositivity defined by IC_50_ values above the assay lower cutoff. Y axis: 0-50% seropositivity.

Seropositivity patterns were broadly consistent across the three cohorts, with the highest titres observed against XBB.1.5 PVs, followed by BA.2.75 PVs (Figure 3a). In contrast, seroprevalence to BA.2.86 and JN.1 PVs was markedly lower in spring 2025, while differences among variants were less pronounced in spring and autumn 2024, particularly relative to BA.2.75 and XBB.1.5 PVs. Neutralising antibodies to the ancestral Wuhan PV remained consistently the least prevalent across all cohorts (Figure 3a).

**Figure 3.**
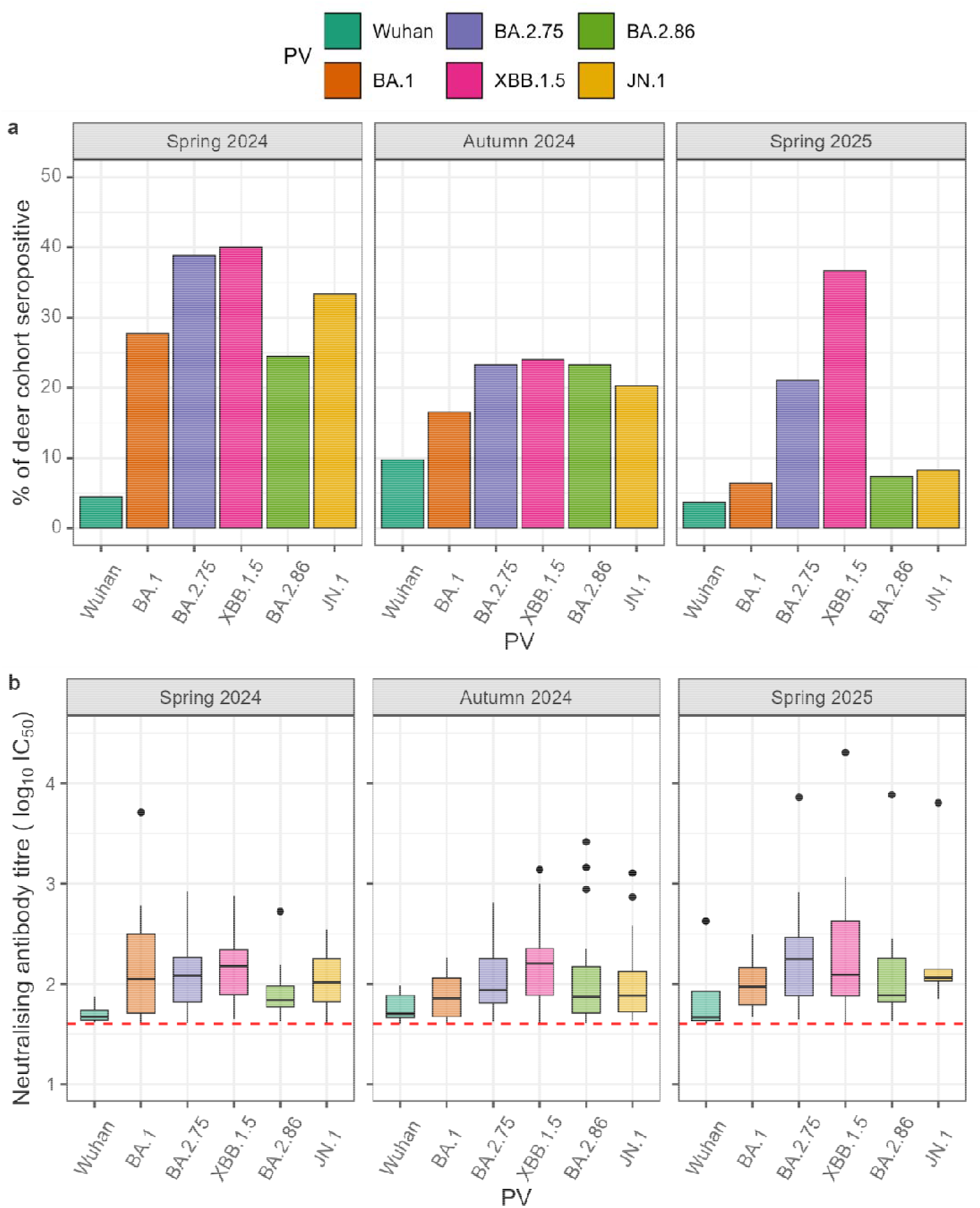
a) Seroprevalence of positive neutralising antibody responses to a wide range of SARS-CoV-2 spike variant PVs, spanning 2019 (Wuhan) to 2025 (JN.1). Responses stratified by each cull cohort: spring 2024, autumn 2024 and spring 2025. Seropositivity defined by IC_50_ values above the assay lower cutoff. Y axis: 0-50% seropositivity. b) Neutralising antibody titres (log_10_IC_50_) against SARS-CoV-2 spike PVs in deer sera culled in spring 2024, autumn 2024, and spring 2025. Only positive titres are shown. Boxplots display the median, interquartile range (IQR), and distribution of titres for each variant (Wuhan, BA.1, BA.2.75, XBB.1.5, BA.2.86 and JN.1). Outliers are further than 1.5 times the IQR from the upper and/or lower whiskers, respectively, and are plotted individually. Assay lower cutoff denoted by a red dashed line.

Seroprevalence of neutralising antibodies against BA.1, BA.2.75, BA.2.86 and JN.1 spike-bearing PVs decreased across each successive cohort (Figure 3a). However, XBB.1.5 PV-neutralising antibodies decreased in prevalence from spring to autumn 2024, before increasing again in spring 2025 (Figure 3a). Despite this increase, median neutralising antibody titres for XBB.1.5 spike PV did not increase from autumn 2024 to spring 2025 (Figure 3b). Similarly, reductions in seroprevalence for other PVs were not consistently associated with lower neutralising antibody titres (Figure 3b). Neutralising antibody titres to the Wuhan spike-containing PVs were consistently lower than those of Omicron-lineage across all three cohorts (Figure 3b).

### Distribution based on sex of deer

Of the 332 deer samples, 59% (197/332) of were male and 41% (135/332) were female deer. The spring 2024 and spring 2025 cohorts were entirely (100%) or predominantly (94%) male deer, respectively. Female deer predominated in the autumn 2024 cohort (97%; Table 2). Seropositivity rates in male and female deer were 43% (85/197) and 42% (57/135), respectively, showing no evidence of sex bias (Figure 4a). Similarly, male and female neutralising antibody titres were comparable, with slightly higher titres in male deer (median titre = 1.99, *n* = 243) than in female deer (median titre = 1.92, *n* = 156; Figure 4b).

**Figure 4.**
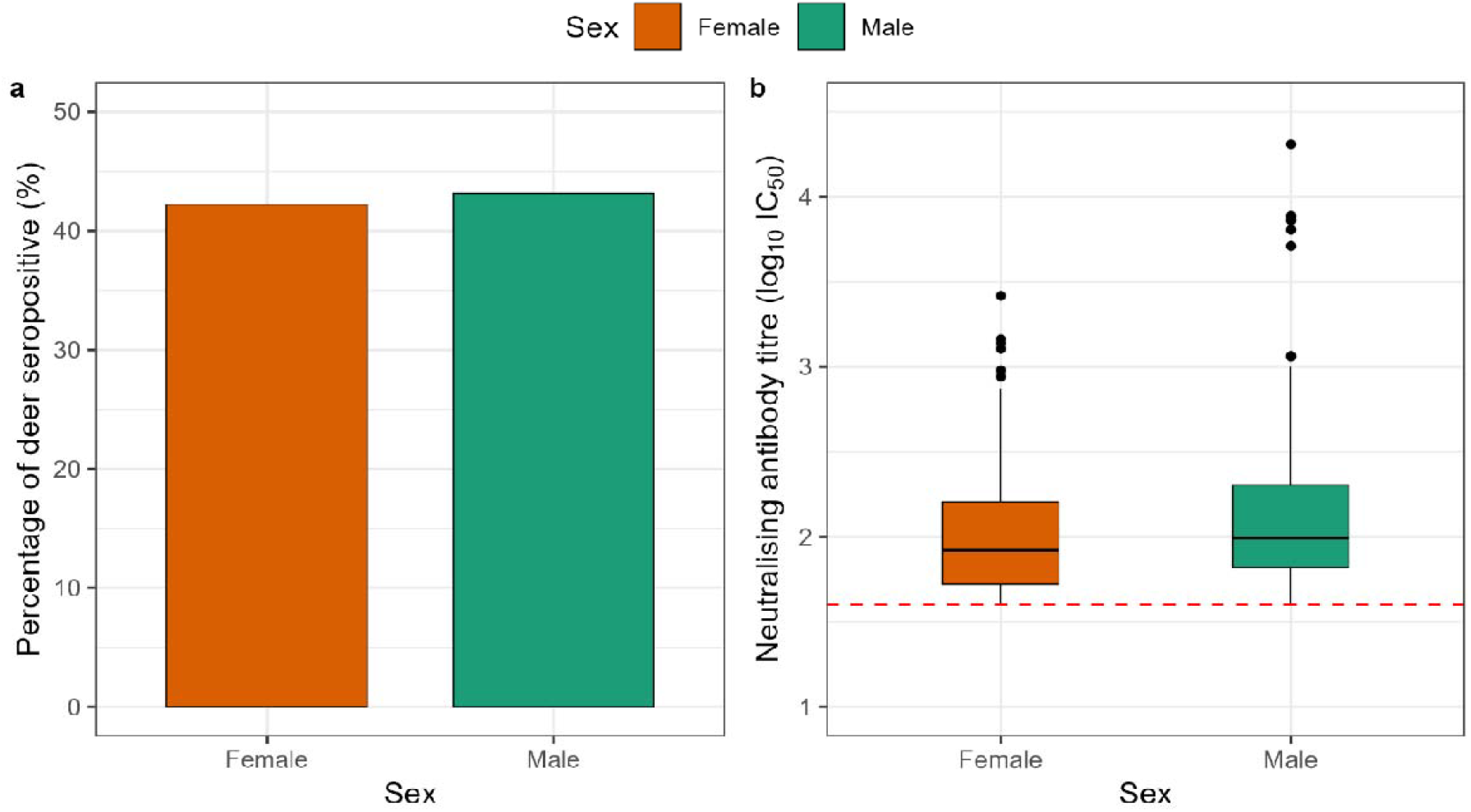
a) Seroprevalence of positive neutralising antibody responses to at least one SARS-CoV-2 spike variant PV. Responses stratified by sex. Seropositivity defined by IC_50_ values above the assay lower cutoff. Y axis: 0-50% seropositivity. b) Neutralising antibody titres (log_10_IC_50_) against SARS-CoV-2 spike PVs in deer sera, stratified by sex. Only positive titres are shown. Boxplots display the median, interquartile range (IQR), and distribution of titres for each variant (Wuhan, BA.1, BA.2.75, XBB.1.5, BA.2.86 and JN.1). Outliers are further than 1.5 times the IQR from the upper and/or lower whiskers, respectively, and are plotted individually. Assay lower cutoff denoted by a red dashed line.

### Species differences

62% (98/158) of fallow deer sera showed a neutralising antibody response to at least one SARS-CoV-2 spike PV, compared to 25% (44/174) of red deer samples (Figure 5a). This species disparity was seen across all PVs tested, for example 56% (88/158) of fallow deer were seropositive to XBB.1.5 spike PV compared to 11% (20/174) of red deer (Figure 5b). Both fallow and red deer exhibited higher seroprevalence to the XBB.1.5 spike compared to more the contemporary BA.2.86 and JN.1, reflecting overall trends (Figure 1). Neutralising antibody titres were higher in fallow deer (median titre = 2.03, *n* = 318) than in red deer (median titre = 1.80, *n* = 80; Figure 5c).

**Figure 5.**
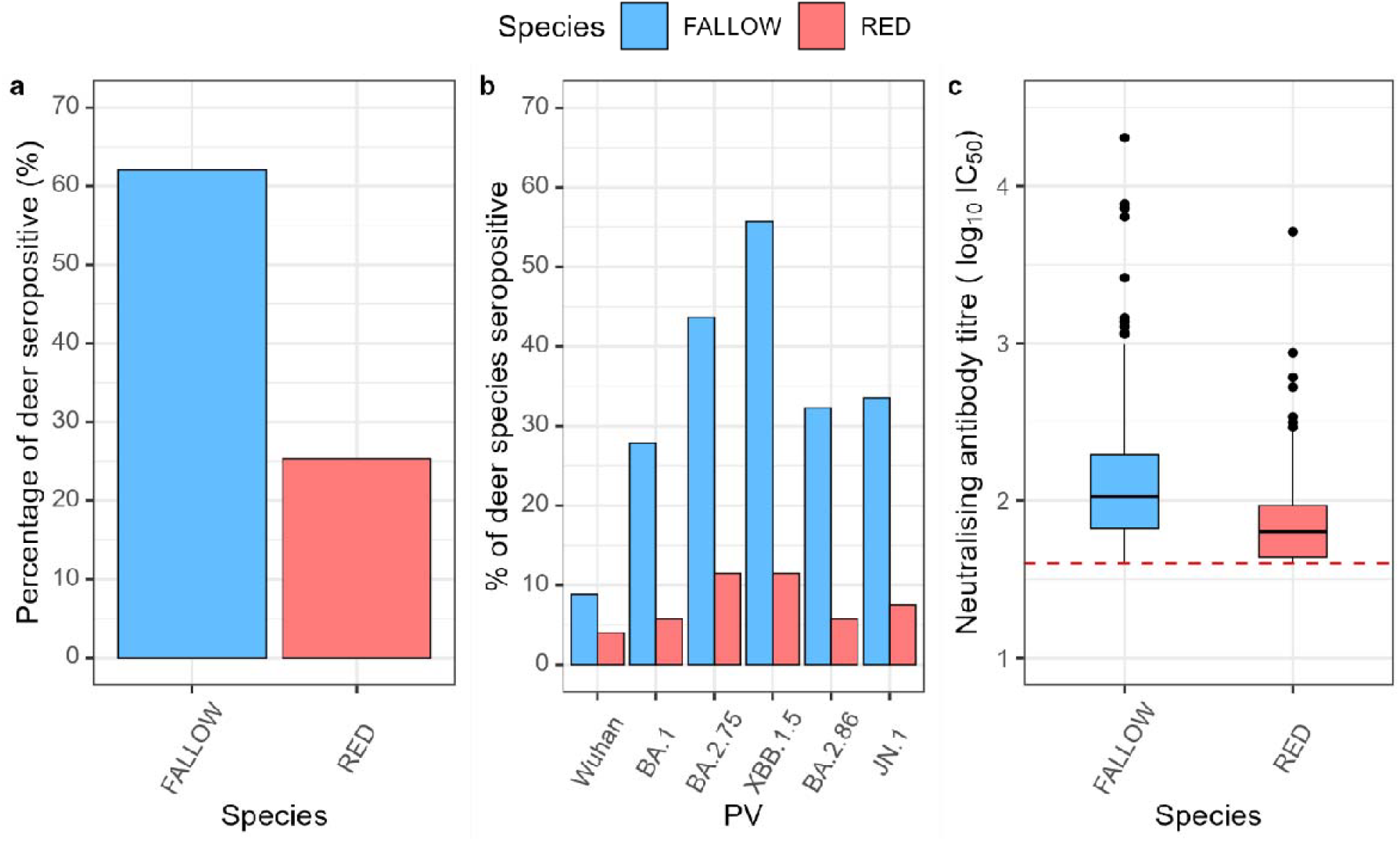
a) Seroprevalence of neutralising antibody titres to at least one SARS-CoV-2 spike PVs, stratified by species: fallow deer and red deer. b) Seroprevalence of neutralising antibody titres to each SARS-CoV-2 spike PV tested, stratified by fallow deer and red deer. Seropositivity in all figures defined by IC_50_ values above the assay lower cutoff. c) Neutralising antibody titres (log_10_IC_50_) against all SARS-CoV-2 spike PVs, stratified by species: fallow deer and red deer. Only positive titres are shown. Boxplots display the median, interquartile range (IQR), and distribution of titres for each variant (Wuhan, BA.1, BA.2.75, XBB.1.5, BA.2.86, and JN.1). Outliers are further than 1.5 times the IQR from the upper and/or lower whiskers, respectively, and are plotted individually. Assay lower cutoff denoted by a red dashed line.

### Age

In spring 2024, the highest seroprevalence of neutralising antibodies was observed in the 2Y age group (Figure 6a). However, by autumn 2024 and spring 2025, the peak seroprevalence had shifted into the 4Y and >5Y age groups, respectively. The median age of seropositive deer was 2Y in spring 2024, and 4Y in autumn 2024 and spring 2025. 82% (9/11) of >5Y deer in spring 2025 were seropositive to at least one SARS-CoV-2 spike variant PV, but these responses were generally low titre (Figures 6a and 6b).

**Figure 6.**
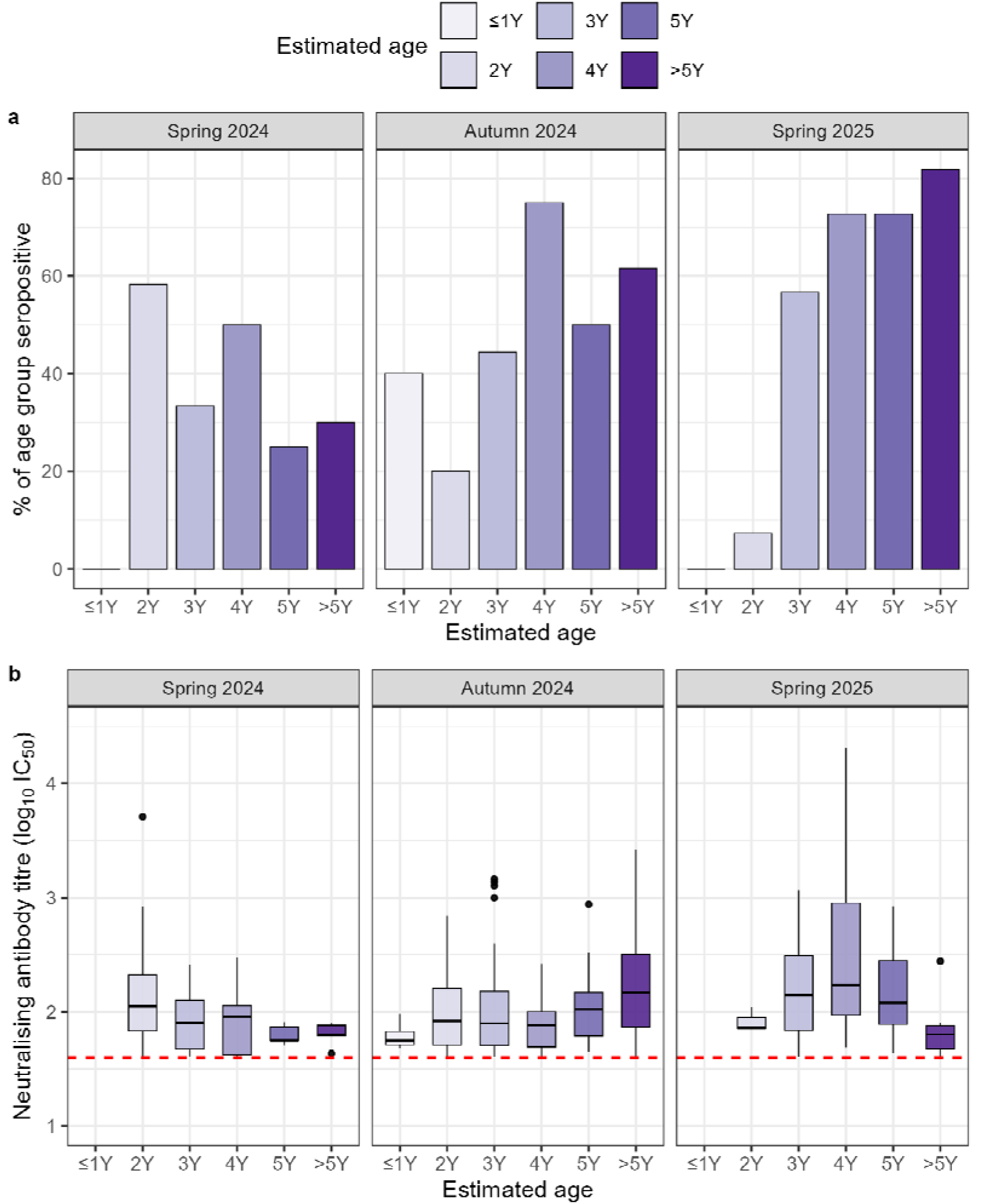
a) Seroprevalence of neutralising antibody titres to SARS-CoV-2 spike PV variants (Wuhan, BA.1, BA.2.75, XBB.1.5, BA.2.86, JN.1). Responses stratified by age: less than 1 year old (≤1Y), 2 years old (2Y), 3 years old (3Y), 4 years old (4Y), 5 years old (5Y) and more than 5 years old (>5Y). Seropositivity defined by IC_50_ values above the assay lower cutoff. b) Neutralising antibody titres (log_10_IC_50_) against SARS-CoV-2 spike PVs in deer culled in spring 2024, autumn 2024, and spring 2025, stratified by age: less than 2 years old (<2Y), 2 years old (2Y), 3 years old (3Y), 4 years old (4Y), 5 years old (5Y) and more than 5 years old (>5Y). Only positive titres are shown. Boxplots display the median, interquartile range (IQR), and distribution of titres for each spike variant (Wuhan, BA.1, BA.2.75, XBB.1.5, BA.2.86, JN.1). Outliers are further than 1.5 times the IQR from the upper and/or lower whiskers, respectively, and are plotted individually. Assay lower cutoff denoted by a red dashed line.

Similarly, the highest neutralising antibody titres were detected in the 2Y age group from spring 2024, while the >5Y and 4Y age groups showed the highest median titres in autumn 2024 and spring 2025, respectively (Figure 6b). Overall, there is a shift in seroprevalence and neutralising antibody titres towards older age groups through successive sampling cohorts, likely reflecting deer historically exposed to SARS-CoV-2 which are aging through the population and subsequent culls.

Seropositivity is zero or low in the ≤1Y and 2Y groups from spring 2025, which includes responses to more recently emerging strains, such as BA.2.86 and JN.1 (Figure 6a). The only seropositive antibody responses seen in the ≤1Y age group were from two 1 month old calves in the autumn 2024 cohort. These responses had low neutralising antibody titres and likely represent maternally-derived antibodies (Figure 6b). The trend of decreasing seropositivity observed in younger age groups could indicate that newer generations of deer have not been infected with contemporary strains.

## DISCUSSION

SARS-CoV-2 is known to infect and replicate in a wide range of mammals associated with humans and their environment. This broad host tropism indicates that there are many reservoir species that could possibly sustain viral populations that could spill back into human populations. The data presented from an urban British deer park with high human footfall. There are many such parks in Great Britain that were heavily visited during the COVID-19 pandemic, especially for daily exercise, one of the few permitted forms of outdoor activity during the ‘lockdown’ periods^36^. These parks continue to be used recreationally, potentially enabling deer to encounter contemporary viral variants of SARS-CoV-2 from infected humans.

Here, we provide strong serological evidence of infection in fallow and red deer species found in Great Britain, as measured by the presence of neutralising antibodies. Seroprevalence of neutralising antibodies were highest against the Omicron-lineage spike-containing PVs, reflecting variants which circulated during 2021 – 2025 (BA.1, BA.2.75, XBB.1.5, BA.2.86 and JN.1) and exhibited little response against the Wuhan spike from 2019/2020 (Figures 1, 3, 5b). Low seropositivity and antibody titres for Wuhan spike-containing PVs may be due to antibody waning since first challenge in 2019/2020, or loss of seropositive individuals from the population over time (i.e. routine culling or natural death). The higher seroprevalence of neutralising antibodies to Omicron-lineage variants may reflect more recent infections, particularly XBB.1.5, BA.2.86 and JN.1 which circulated in 2023 and 2024/5, the individuals of which have yet to undergo multiple rounds of culling.

In spring 2024, 2 year old deer showed the highest rates of seropositivity to SARS-CoV-2 PVs, compared to other age groups (Figure 7). By spring 2025, this had shifted to the older age groups displaying highest seropositivity, particularly the >5Y age group. This may indicate that exposure rates are declining, as 2-year-old deer culled in spring 2025 had a lower prevalence of neutralising antibodies than in the previous year. Additionally, the increasing seroprevalence with age is indicative of cumulative exposure to SARS-CoV-2 over time, as the disease has become endemic in human populations.

Behavioural and physiological differences have been hypothesised to bias infectious disease incidence towards male animals and may affect management strategies in wild deer^37,38^. However, seropositivity rates and neutralising antibody titres were comparable between male and female deer, suggesting that sex-related factors had not affected disease transmission in this population.

The data we have presented highlights a clear difference between native British red deer and fallow deer, which seem particularly susceptible - whether immunologically or as a result of behavioural differences - to infection by this virus. Intrinsic differences in species biology may influence permissibility and transmissibility of viral infection, especially ACE2 receptor variations. Tissue tropism has been identified in the lower respiratory tract in both deer species^27^. However, Omicron variants have shown increased replication in the upper respiratory tract^39^. Further study of ACE2 receptor abundance in the upper respiratory tract in red deer and fallow deer may help to explain our observed species differences. Secondly, characterisation of ACE2 receptor structure and binding affinity will help to highlight species variability which could have resulted in increased susceptibility in fallow deer. Importantly, this data covers only one managed deer park among dozens in the UK, and only two out of six distinct species currently found in Great Britain, some of which are introduced species from multiple worldwide origins. Extending this work to the other four species of GB resident deer would add valuable insight into the viral reservoir capacity provided by deer in the country, whether in parks or wild populations.

Behavioural considerations affecting the rate of contact with humans or other deer may account for the disparity in seropositivity observed between red and fallow deer. Factors such as living in large social groups, occupying high-traffic areas of the park, and being more habituated to humans may increase infection rates, either through more frequent human-deer contact or increased deer-to-deer transmission. The underlying mechanisms are unclear and likely multifactorial. Additionally, species variation in behaviour, such as differences in home range size, may have further impacts on disease spread beyond individual deer populations. Future research evaluating human-deer and deer-deer interactions, both in managed parks and in the wild, could help contextualise these species differences, inform risk assessments, and guide strategies to protect both public and wildlife health.

## Limitations

Our samples were collected during routine culling, which is likely to affect the seroprevalence dynamics within this closed population. A proportion of seropositive individuals will have been removed from the population and replaced by naïve calves during the birthing season, which may contribute to the reduced seroprevalence of neutralising antibodies in the spring 2025 cohort compared to the original spring 2024 cohort despite no significant decrease in antibody titres.

In white-tailed deer in the United States, SARS-CoV-2 neutralising antibodies have been shown to persist for at least 13 months^24^. However, antibody titres may wane over time or be boosted if challenged with homologous and cross-reactive strains. Without any further data on antibody kinetics in red and fallow deer, it remains difficult to infer infection history from antibody titres alone, such as when they were infected or exactly which strain(s) they were infected by. Whole genome sequencing data could be used to identify infective strains and elucidate transmission dynamics between human and deer in cases where viral RNA has been detected, as illustrated in other studies of white-tailed deer in the USA^18,23,40^.

There could also be differences between antibody kinetics in young and old deer, for example more rapid antibody waning in older deer, which makes robust comparisons between age groups difficult at this stage. In each cohort, the 2Y age group predominates, which is not representative of the wider deer population. Furthermore, our methods do not allow for longitudinal measurements of antibody titres over time within individuals. This would allow us to track antibody waning or identify increases in titres which may be suggestive of a recent exposure during the sampling period.

Each cull cohort was skewed towards a particular sex: spring 2024 and spring 2025 cohorts were predominantly male, whereas the autumn 2024 cull cohort was mostly female. Sex-based behavioural differences may influence exposure dynamics and thus neutralising antibody prevalence. Comparisons between the female-biased autumn 2024 cohort and the male-biased spring 2024/2025 cohorts should therefore be interpreted with caution. For example, the apparent increase in XBB.1.5 PV seropositivity from autumn 2024 to spring 2025 may be confounded by sex differences. Male and female deer were not equally represented overall across the 332 samples.

SARS-CoV-2 is part of the subgenus *Sarbecovirus*, within the *Betacoronavirus* genus, distinguishing these viruses from the related seasonal coronaviruses, HCoV-OC43 and HCoV-HKU1, which are embecoviruses. The other seasonal coronaviruses HCoV-229E and HCoV-NL63 are part of the *Alphacoronavirus* genus, further differentiating these viruses from SARS-CoV-2. Therefore, it is unlikely that any previous exposure to seasonal coronaviruses would result in cross-reactive neutralising antibodies detectable in our SARS-CoV-2 spike PV assay^41^.

Further assays, such as enzyme-linked immunosorbent assay (ELISA) for SARS-CoV-2 nucleocapsid (N) protein could be used to increase the specificity of our findings, by confirming binding of antibodies to an internal SARS-CoV-2 protein. Additionally, live virus neutralisation assays could be used to confirm specific neutralisation, although a high level of correlation between pseudotype and live virus neutralisation has been reported^42^.

## Conclusions

Interactions between humans and deer populations are likely to become more frequent through continual urban expansion and changing behaviour in wildlife, leading to increased opportunities for zoonotic pathogen transmission. The COVID-19 pandemic initially occurred as a result of high-risk human behaviour relating to wildlife species (live animal markets) that can harbour viruses of human relevance^43^. Despite clear guidance in these managed parks that wildlife should not be approached and should be observed at a distance, our findings show serological evidence of SARS-CoV-2 infection. With no reported increase in morbidity or mortality, it appears that SARS-CoV-2 infection is generally subclinical in these deer populations and does not pose a threat to herd health. However, this does not preclude the possibility of these deer acting as a reservoir in which the virus can circulate and evolve, which may then be transmitted back into human populations.

Wildlife populations pose a unique challenge for zoonotic disease emergence, with relatively little monitoring and surveillance, but close proximity to humans in many urban and peri-urban areas. SARS-CoV-2 exposure of different deer species in Great Britain, the Republic of Ireland, Canada and the United States are clear evidence that these animals could represent an overlooked reservoir for the virus. Surveillance is integral to safe and sustainable coexistence with wildlife species, to promote preparedness and safeguard both human and animal health from endemic and emerging zoonotic viruses.

## MATERIALS AND METHODS

### Blood sampling

Bloods were collected from animals already destined to be culled for population management and the venison market. 8ml of whole blood was taken at the point cadavers were gralloched and kept at 4°C until processing into serum. Serum aliquots were made and heat inactivated at 56°C for 1h before use in the below immunoassays.

### Cells and viruses

HEK293T/17 cells were obtained from ATCC (CRL-11268 ™)

### Pseudotyped lentivirus production

Spike-bearing lentiviral pseudotypes were produced by transient transfection of HEK293T/17 cells with packaging plasmids p8.91 and pCSFLW and different spike-bearing expression plasmids in the pcDNA3.1 plasmid backbone, using the Fugene-HD (Promega, E2311) transfection reagent. Supernatants were harvested after 48⍰hours, passed through a 0.45 µm cellulose acetate filter, and titrated on HEK293T/17 cells transiently expressing human ACE-2, and TMPRSS2. Target HEK293T/17 cells were transfected, 24⍰hours prior to infection, with 2⍰µg pCAGGS-huACE-2 and 150⍰ng pCAGGS-TMPRSS2 in a T75 tissue culture flask.

### Pseudotype based microneutralisation assay

Pseudotype-based micro-neutralisation assays (pMN) were performed as described previously^35^. Serial dilutions of serum were incubated with lentiviral pseudotypes bearing sarbecovirus spike proteins for 1⍰hour at 37°C, 5% CO_2_ in 96-well white cell culture plates. 1.5 × 10^4^ HEK293T/17 cells transiently expressing human ACE-2 or DPP4 and TMPRSS2 were then added per well and plates incubated for 48⍰hours at 37°C, 5% CO_2_ in a humidified incubator. Bright-Glo (Promega) was then added to each well and luminescence read after a five-minute incubation period. Experimental data points were normalised to 100% and 0% neutralisation using cell only and PV-only cell control wells respectively and non-linear regression analysis performed in GraphPad Prism 10.4 to produce neutralisation curves and resulting IC_50_ values. NIBSC antibody standards were employed as positive controls in all assays. Sera were tested starting at a 1:40 dilution, giving the assay a lower sensitivity limit of Log_10_(40). Seropositivity was therefore defined as an IC_50_ value above this assay cut-off.

### Data visualisation

Data visualisation was performed in RStudio (R version 4.5.0)^44^. Packages used: readxl(), tidyr(), ggplot2(), dplyr(), ggpubr(), for data manipulation and visualisation^45–49^.

## Ethics

Ethical approval was obtained under the non-regulated route by the Animal Welfare and Ethical Review Board (AWERB), University of Cambridge.

## Acknowledgments

The authors thank Kei Sato for the JN.1 expression plasmid, and Rachael Tarlinton, Steve Dunham and Leah Goulding for useful discussion in deer and wildlife virology.

## Data and materials availability

All data are available in the manuscript or at reasonable request from the corresponding author.

